# AuthormetriX: Automated Calculation of Individual Authors’ Non-Inflationary Credit-Allocation Schemas’ and Collaboration Metrics from a Scopus Corpus

**DOI:** 10.1101/2025.01.19.633820

**Authors:** Samuel O. Adeosun

## Abstract

**Background:** Publication count is the currency in academia, but the most widely used whole count method is considered unfair and inflationary. Several non-inflationary author credit-allocation schemas (NIACAS) have been proposed, but none is widely adopted in practice (e.g., in evaluating faculty scholarly productivity) or in bibliometric research. This low adoption and implementation rate may be due to the complexity in operationalizing the schemas.

**Aim:** To develop an application that automates the calculation of individual authors’ scholarly output metrics based on multiple NIACAS.

**Method:** Published formulas of NIACAS were written as Python functions wrapped in a Streamlit user interface that takes .*csv* files of the relevant corpus, and a list of author’s Scopus IDs as inputs. The functions calculate the authors’ output metrics based on NIACAS including first- and last-author straight counts, arithmetic, golden share, and multiple variations of fractional, geometric, and harmonic schemas. Collaboration metrics are also calculated. Secondary features include modelers for author counts and schemas. In a use-case, absolute rank displacement (ARD) and actual contribution proportions (ACP) were compared between highly cited clinical medicine researchers and high *h*-index pharmacy practice faculty populations.

**Results:** AuthormetriX accurately calculates individual authors’ aggregate scholarly output based on 14 NIACAS, from the file inputs. There were schema and population differences in ACP, but only schema differences in ARD within the populations studied.

**Conclusions:** AuthormetriX simplifies the implementation of non-inflationary author credit-allocation schemas and will facilitate their broader adoption in practice and bibliometric research. AuthormetriX is freely available at https://sadeosun-a-uthormetrix-v1.streamlit.app/.

## 1.0 INTRODUCTION

Scholarly publishing is a key job expectation in academia. The “Publish or Perish” aphorism in academia accurately represents the fact that publication count is a proxy for scholarly productivity,[1] and the “currency of the realm.”[2,3] Publication counts are used in evaluating faculty for career-advancing benefits including faculty positions, grants, tenure, promotions, and awards.[2,4,5] Other ‘assets’ like high *h*-index, making the highly cited researchers list, and reputation as an expert in the field are driven by publication counts.[2,3,6,7] This reward system incentivizes faculty to *be an author* on as many publications as possible. This is because these academic assets are based on the whole count method which gives full credit to every author on a document regardless of the number of co-authors, thus making an individual’s single- and multi-author publications weigh the same in their publication counts.[1,8] Consequently, single-author publication rates have seen drastic declines while the proportion of multi-author publications, and the number of authors per publication have steadily increased in every discipline.[9–11] Derek de Solla Price and others noticed this multi-authorship trend and its implications as early as the 1960s[12] and said: “…*part of the social function of collaboration is that it is a method for squeezing papers out of the rather large population of people who have less than a whole paper in them*.” Furthermore, he considered that using the whole count method makes multi-author publications “*a very cheap way of increasing apparent productivity*.*”*[13] Therefore, to address this inflation in publication credits in the academic economy, Price and others suggested that the whole count method was no longer reasonable since single-author publications were no longer the norm. They suggested that the credit from multi-author publications must be divided *equally* among all the co-authors. This credit schema was called adjusted, or fractional credit allocation.[12– 14]

While the fractional credit allocation schema addresses the inflationary bias in the whole count schema,[14–17] others have pointed out the unfairness, and yet another bias (equalizing bias) in the fractional credit-allocation schema due to the assumption that all authors contributed equally.[16–19] Hagen, stated that equalizing bias introduces “*a large counterproductive element of reverse meritocracy and systemic injustice into bibliometric performance assessment*.*”*[20] Thus, other schemas allocate credits to authors based on their positions on the authorship byline. This sequence-determines-credit (SDC) convention was recognized as early as the 1960s[21] and has been supported by studies showing that the author sequence represents their relative contributions to the publication.[22–24] The exception to this convention is in fields where alphabetic ordering is still used.[10] Therefore, based on the SDC convention, first authors are known to have done the bulk of the work,[22–25] which is the basis of the straight first author method, where the whole credit goes to *only* the first author.[25,26] Conversely, in recognition of the leading roles and major contributions of senior or corresponding authors who are often listed last on the authorship byline,[23,27,28] some schemas recognize only the last author (straight last author).[29,30] Other schema variants emphasize both authors (first and last author emphasis; FLAE e.g., harmonic FLAE[31]; Fractional FLAE,[32]) as they are considered the most important contributors compared to middle authors.[11,22–24] The non-inflationary credit allocation schemas (NIACAS) and variants included in this study are shown in Table 1. These NIACAS (Schemas 2-15) follow the general principles that address the inflationary bias in the whole count method (Schema 1). Therefore, they meet the “strictly non-inflationary” rule (i.e., total credit must be 1 for a single document),[33] independent of the first and third of the 3 theoretical and practical principles that credit allocation schemas are supposed to meet. The principles include[15,26]:

**Table 1.**
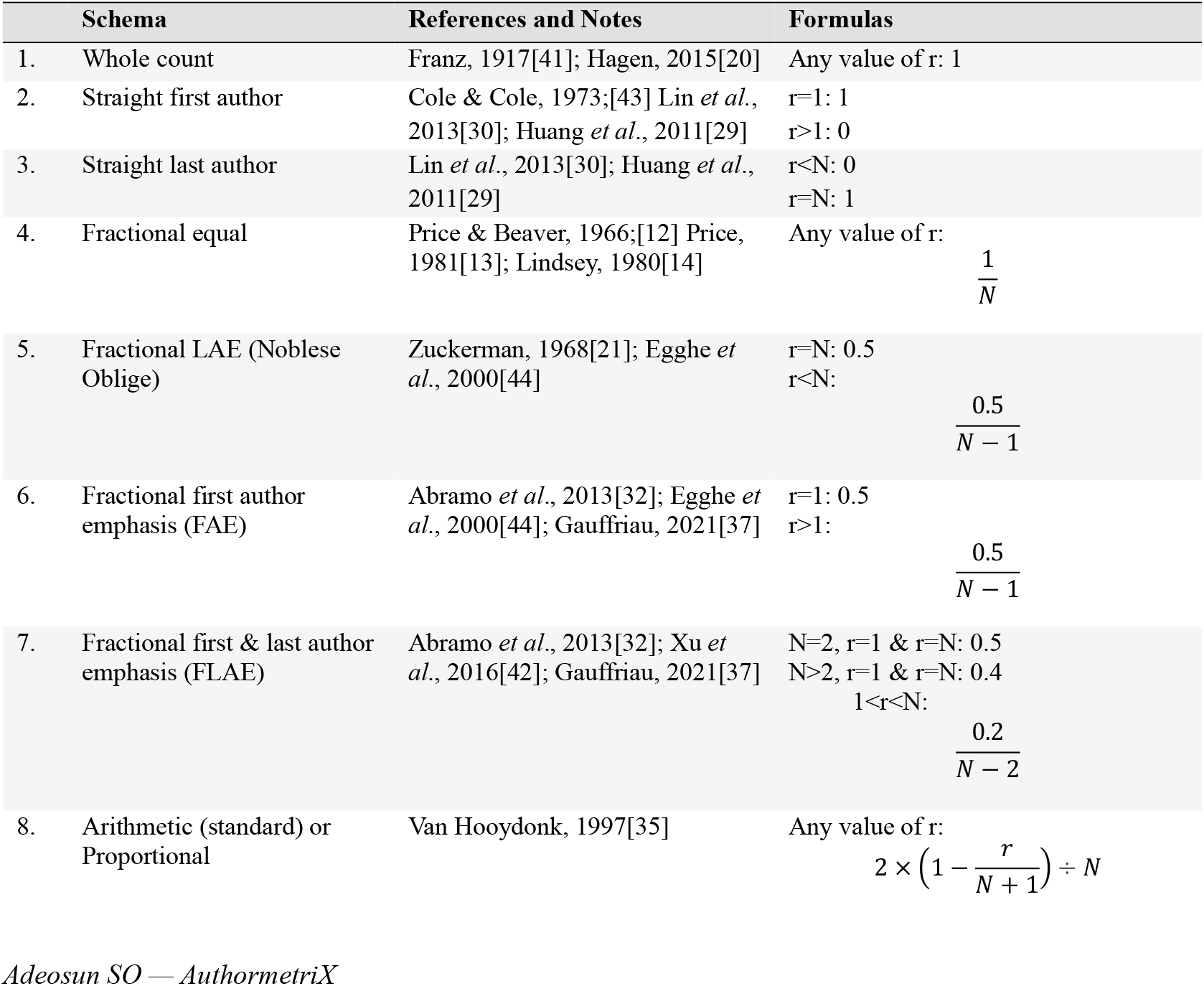
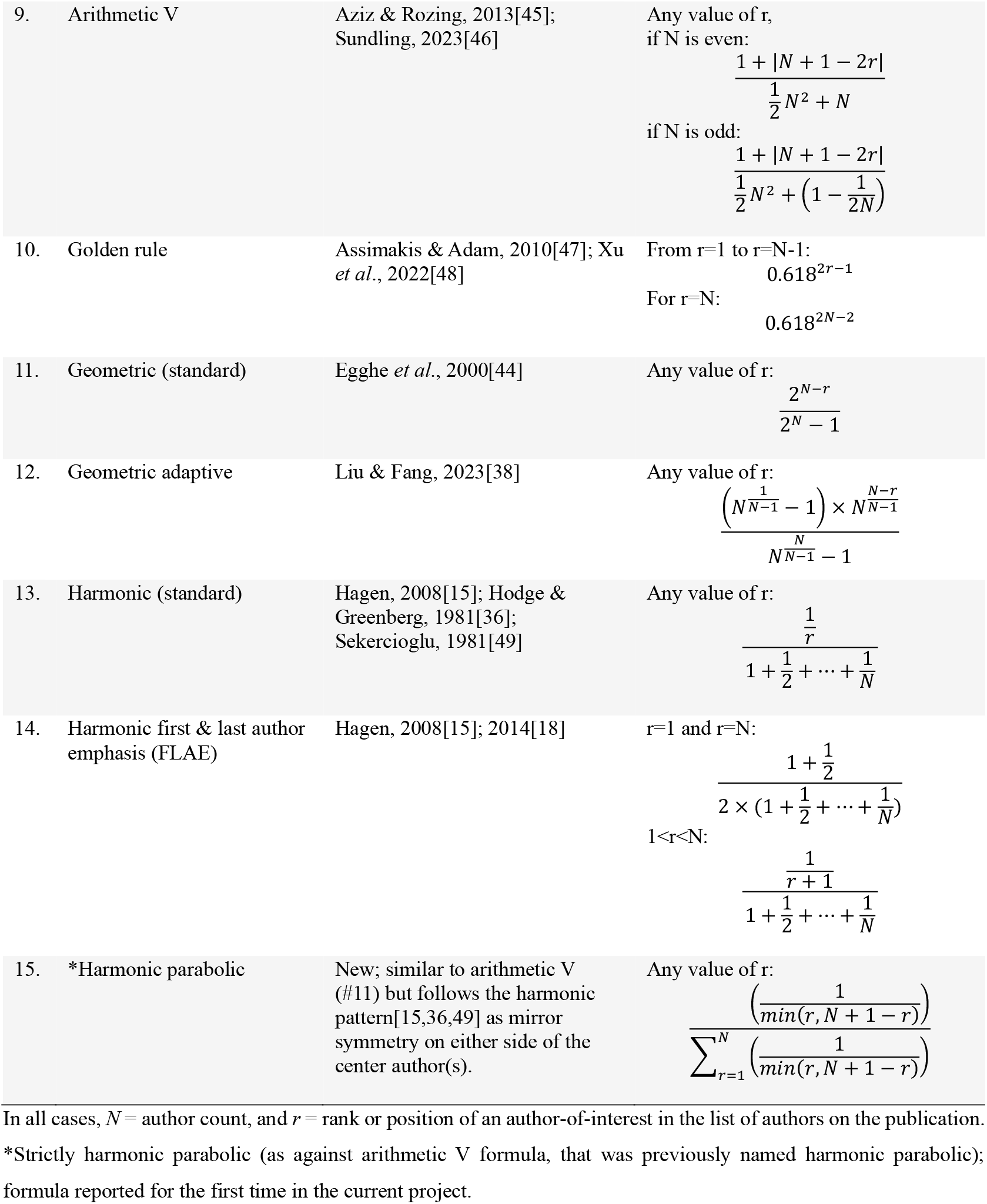
Schemas and Formulas Used in the Python Functions in AuthormetriX.

a. Total publication credit is shared among all coauthors,
b. The first author gets the most credit, the *r*^th^ author receives more credit than the (*r+1*)^th^ author, and
c. The larger the number of authors (author count), the lower the credit per author.

Despite the consensus on the inflationary bias in the whole count schema and the equalizing bias that remains in fractional counting,[13,14,16,18,31,34,35] none of the schemas proposed to address the biases have seen widespread adoption.[15,17,36,37] This is likely due to the complexity in these formulas (despite having only 2 parameters; author count (*N*), and position of an author-of-interest (*r*)),[38–40] and the prohibitive, labor-intensive process involved[20] compared to the whole count method. Despite the inherent flaws, the simplicity and ease of implementing the whole count method had sustained its dominance since its first use by Franz in 1917.[20,41] Therefore, the aim of this project is to build an application that simplifies the calculation of several NIACAS, to leverage the inherent fairness and validity for implementation in practice (e.g., award or tenure and promotion committees’ evaluations, etc.) and in bibliometric research.

## 2.0 METHODS

Existing reviews with[42] or without formulas[37] were reviewed first. Additional literature reviewed either proposed a new NIACAS, or further described previously proposed schemas.[12–15,18,20,21,29,30,32,35– 38,41–49] The current study focused on schemas with all author credits summing up to 1.0, and with no additional parameter required beyond the total number of authors (author counts; *N*), and the rank (*r*) of the author(s)-of-interest in calculating authors’ credits. Table 1 shows the schemas, formulas, primary and/or secondary supporting literature, and notes. After writing Python functions based on each formula, the unnamed formula originally proposed by Aziz & Rozing,[45] which was later named “harmonic parabolic”,[46] was found to be neither harmonic in mathematical terms, nor parabolic in shape. Therefore, that schema was referred to as arithmetic V in this study (according to the mathematical model and shape of the plot). A Python function with a logic that follows the harmonic model with a mirror-symmetry on either side of the center author(s), (i.e. a parabolic plot shape) was written. With some assistance of DeepSeek-V3,[50] the function was translated to a simple mathematical formula similar to the standard harmonic formula[15,36,49] requiring the same 2 variables (author count, and author-of-interest’s position). The function was then re-written based on the formula, to further simplify the code. All codes were written in Python (version 3.10.11) in Visual Studio Code (VSCode version 1.96.3). Code completion assistance was provided by Codeium (version 1.30.18) –a VSCode extension. Code completion suggestions were only accepted after deeming them correct. Python libraries used were

### 2.1 Automated Calculation of Individual Author Credits and Collaboration Metrics

The corpus supplied is pre-processed by first removing rows (documents) with missing information in any of the columns: ‘Author(s) ID’, ‘Document Type’, and ‘Year’, and then deduplicated based on ‘Author(s) ID’, ‘Title’, ‘Source title’, and ‘Year’ based on previous experience with analyzing Scopus data.[9,51].

Each NIACAS formula written as individual Python sub-function takes the author count (*N*) as the only argument and returns a list of credit-allocation of the same length as the number of authors in each document. The mother function contains a line to apply each sub-function to the author count of each document in the corpus. The output of that line (a list of the credits allocated to each author according to the author order) is a new column in the corpus named according to each NIACAS.

The second user-supplied dataframe with the Scopus ID (SCID) of each author-of-interest (one per row, required to be in the first column) is de-duplicated. This dataframe and the pre-processed corpus dataframe then pass into functions that search for each SCID-of-interest in each document’s Authors’ ID list. For each NIACAS, the credit that corresponds to the position of the SCID in the Author IDs list of each document is aggregated, and the sum of those credits is returned to the respective row (of SCID) under the relevant NAICAS column in the SCIDs dataframe.

The 2 dataframes then pass on to other functions that return collaboration metrics, and other related metrics for each author-of-interest. These metrics include the Degree of Collaboration (DC),[52] Collaboration Index (CI, based on multi-author publications only),[53,54] Collaboration Coefficient (CC),[55] number of single-author publications, and number of unique co-authors (*not* including the author-of-interest him/herself). Another metric calculated is the first and last authorship proportion (FLAP), based on the premise that the first and last authors are the major contributors to a publication.[22–24] FLAP is the ratio of the sum of straight first, and straight last author counts, to the whole count of an individual. The final output dataframe is then displayed for the user to download.

To further verify the accuracy of all codes and functions in the application, manual calculations of the metrics were done (with Microsoft Excel), using an author with 15 publications, in which the author occupied a variety of ranks (first, last and middle), among 2 to 12 total authors of those publications.

#### 2.1.1 User Interface and Processing Steps

The user interface was built on the Streamlit framework. The analysis process involves only 3 total steps (which includes 1 optional step). The first step involves uploading a .*csv* file of the relevant corpus. This is followed by the optional step where the uploaded corpus can be filtered by either one or both of ‘document type’, and the ‘publication year’. The last step involves uploading the .*csv* file containing the

SCIDs-of-interest (i.e., Scopus IDs of the authors-of-interest). The home page includes short video how-to guides for obtaining the two input files needed. The final output is the downloadable SCIDs table updated with columns for all 21 scholarly output and collaboration metrics calculated.

### 2.2 Author Count and Schema Modeling

The secondary features include 2 modelers. The schema modeler takes a single argument (the total number of authors, *N*), and returns a dataframe of all author positions, and the respective credits based on the 12 NIACAS that allocates non-zero credits to all author positions (i.e., schemas #4-15; Table 1). The Author count modeler takes 2 *different* author count values and returns a dataframe with all author positions and the respective credits allocated based on a single, user-selected schema. For the specified schema, the second modeler highlights how the credits allocated to authors in positions common to both author count scenarios differ. In both modelers, the outputs are available as both a table, and a dynamic visualization in a Plotly line chart.

### 2.3 Example Use Case

To demonstrate how AuthormetriX could be used, a question that requires calculating several NIACAS simultaneously was addressed. The question is based on previous studies showing that the ranks of individuals or other units of analysis (e.g. countries) would be different for different methods used in counting their publications.[18,30,44,45,56]

#### 2.3.1 Population studied

A sample of 50 individuals on the publicly available Clarivate’s 2024 highly cited researchers (HCR) list (Award category = Clinical Medicine; Region = United States)[57] was selected. Using their names and affiliations, their profiles and Scopus IDs were obtained from Scopus.com as was previously described,[51] and in the how-to guide videos in the application home page (https://sadeosun-a-uthormetrix-v1.streamlit.app/). Scopus IDs of Pharmacy Practice faculty from a previous study [51] were sorted by *h*-Index. The top 50 were selected if they had at least 1 article or review document type during 2020-2024. On January 1^st^, 2025, all 50 SCIDS of each population were used to retrieve and download the authors’ publications (corpus), limited by year (2020 through 2024), and Source type (‘Journals’). Using the optional step in AuthormetriX, the publication records were filtered down to articles and reviews only, from which the whole count, the NIACAS, and collaboration metrics were obtained. The outputs were downloaded for further processing in Python.

#### 2.3.2 Outcome measures

The outcome measures used were absolute rank displacement (ARD) and the actual contribution proportion (ACP). The ARD is a slight variation of the (positive or negative) bias-induced rank displacement.[18] For each NIACAS, the ARD is the absolute difference in the ranks of an individual when ranked based on their whole count credits versus each NIACAS credit.

The ACP is the ratio of each NIACAS credit to the whole count credit. NIACAS credits represent sole authorship equivalent[36] which reflects an individual’s total contributions assuming the person published all their *whole* (or *partial*) papers *alone*.[45] The inflation in the authors’ credit, which is also the proportion of credit ascribable to co-authors can be derived from ACP, i.e., 1-ACP. Since ACP is a normalized metric (by whole count), both ACP and 1-ACP are logically comparable across individuals or populations. The 1-ACP value is also consistent with the profit (*p*)-index described as: “*the relative contribution of other individuals to the total publication record of a given author* .”[45] ACP is the inverse of Counting Inflation Ratio (CIR) described by Lin and colleagues.[30] Thus, a lower ACP implies higher CIR and higher *p*-index. The difference between each NIACAS’ ACP and fractional equal ACP also reflects the equalizing bias (positive or negative) detected by the respective NIACAS, as described for the standard harmonic counting schema by Hagen.[18] Positive equalizing bias is detected when fractional equal ACP is greater than a NIACAS ACP, and it means that the bias favored the author, most often at the expense of primary authors, and vice versa when fractional equal ACP is less than a NIACAS ACP.[18]

The ARD and ACP were calculated for each of the 12 NIACAS that strictly meet the first of the 3 principles (i.e., non-zero credits for *all* authors; Schemas #4-15; Table 1). The null hypothesis tested was that the schemas’ ARDs and ACPs of a population of individuals on Clarivate’s highly cited researchers’ lists in the Clinical Medicine award category (HCR Clinical Medicine) are *not* different from those of the high *h*-index Pharmacy Practice (HHI Pharmacy Practice) faculty[51] *within*, and *across* the two populations.

#### 2.3.3 Statistical Analysis

Statistical analysis was done in R software (version 4.4.1).[58] Both outcome measures were modeled using linear mixed effects model with schemas (#4-15; Table 1) and population (HCR Clinical Medicine and HHI Pharmacy Practice) as predictors. The random effect was the individual faculty IDs. ARD was modelled as a Poisson family while ACP was modeled as a beta_family using *glmmTMB*.[59] Model diagnostics was done with the *performance* package.[60] Predictions and pairwise comparisons were made with *ggeffects*,[61] using the Benjamini-Hochberg method to adjust for multiple comparisons. Plots were made with *ggplot2*.[62] Comparisons were done *across* populations *within* each schema. In addition, to determine equalizing bias, within populations, each schema was compared to the corresponding fractional equal. In separate analysis, other metrics were compared across disciplines using independent T-tests. Lastly, the arithmetic V and the [new] harmonic parabolic schema credits were compared within each population (N=50 each), and both populations combined (N=100), using paired T-test. Comparisons with *p*-values <0.05 were considered statistically significant.

## 3.0 RESULTS

AuthormetriX provided accurate results consistent with examples given in the respective publications that proposed the schemas. The same results were also obtained for all metrics calculated manually for the SCID used for verification. However, the manual process was very tedious, time-consuming, and error-prone while AuthormetriX results were obtained in seconds.

In the demonstrated use case, the 50 SCIDs for each of HCR Clinical Medicine and HHI Pharmacy Practice populations retrieved 8274 and 1979 documents respectively. After filtering to articles and reviews in the optional step, there were 7033 and 1738 documents (85.0% and 87.8% of original corpus, respectively) in each corpus for the analyses. The mean (and 95% confidence intervals) *h*-index, for HCR Clinical Medicine and HHI Pharmacy Practice were 113.3 (101.7–124.9) vs. 42.2 (40.6– 43.8), respectively. Other publication- and collaboration-related metrics calculated in AuthormetriX are shown in Table 2. In HCR Clinical Medicine, 13 people (26%) had at least 1 single-author publication versus 8 (16%) in HHI Pharmacy Practice. The total number of those single-author publications were 40 and 24 respectively.

**Table 2.** Other publication and collaboration metrics calculated in AuthormetriX.

|  | HCR Clinical Medicine | HHI Pharmacy Practice |
| --- | --- | --- |
| **Publications** |  |  |
| All (Whole count) * | 158.4 (125.3-191.5) | 35.5 (26.4-44.7) |
| First author* | 8.8 (5.7-11.9) | 3.1 (1.5-4.8) |
| Last author* | 25.6 (19.8-31.3) | 13.6 (9.3-17.9) |
| Single author | 0.8 (0.0-1.6) | 0.5 (0.0-0.9) |
| First or last authorship proportion (FLAP)* | 0.24 (0.20-0.27) | 0.48 (0.41-0.55) |
| **Collaboration** |  |  |
| Degree of collaboration | 0.99 (0.99-1.00) | 0.98 (0.96-1.01) |
| Collaboration coefficient* | 0.90 (0.89-0.91) | 0.82 (0.79-0.85) |
| Collaboration index* | 23.4 (20.1-26.7) | 10.7 (7.8-13.6) |
| Unique co-authors* | 1608.2 (1248.8-1967.6) | 218.5 (129.0-308.0) |

Mean and 95% confidence intervals: \* $p < 0.05$ by independent T-test. Proportions and metrics that range between 0 and 1 are shown with 2 decimal places.

There was no significant schema-population interaction effect in absolute rank displacement (ARD), and each NIACAS ARD was not significantly different across populations (Fig. 1a). However, within each of the two populations, all NIACAS ARD were significantly different from fractional equal (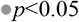Fig. 1a), except arithmetic V and harmonic parabolic (○ *p*>0.05). There was a significant schema-population interaction effect in the actual contribution percentage (ACP; *p*<0.05). While the whole counts (Table 2), and the raw NIACAS credits of HCR Clinical Medicine were higher than those of HHI Pharmacy Practice (Supplementary material ST1), all NIACAS ACP (except golden share and geometric standard) were significantly higher (\**p*<0.05) in HHI Pharmacy Practice than HCR Clinical Medicine (Fig. 1b). Within HCR Clinical Medicine, 7 of the 11 NIACAS were significantly different versus fractional equal. In contrast, within HHI Pharmacy Practice, all 11 NIACAS were significantly different from fractional equal (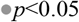 Fig. 1b). The raw arithmetic V and raw harmonic parabolic schema credits were significantly different within HHI Pharmacy Practice (6.8 vs. 6.9; *p*= 0.021) but not within HCR Clinical Medicine (16.7 vs. 16.9; *p*>0.05; Supplementary material ST1), nor within the combined population (11.8 vs. 11.9; *p*>0.05).

**Figure 1.**
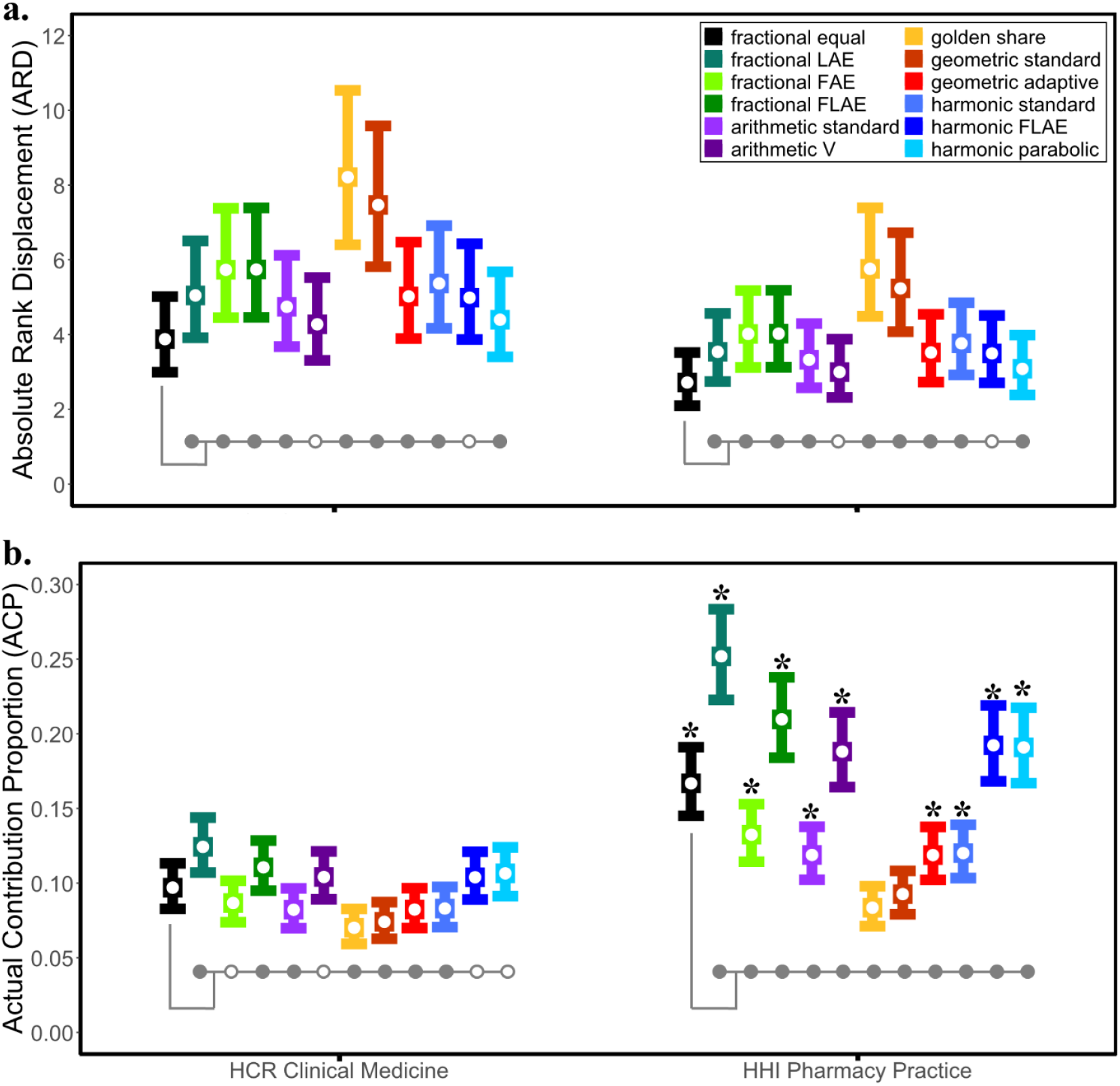
(a) Absolute rank displacement (ARD), and (b) Actual contribution proportion (ACP) based on non-inflationary author-credit allocation schemas (NIACAS). The two populations are samples from the 2024 highly cited researchers (HCR) list Clinical Medicine award category, and from high h-index (HHI) Pharmacy Practice faculty (N = 50 each). Bars represent mean and 95% confidence intervals. \**p*<0.05 (*on HHI Pharmacy Practice) for comparisons of the same schema *across* populations (e.g. HCR Clinical Medicine fractional equal vs. HHI Pharmacy Practice fractional equal). 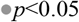(and ○ *p*>0.05) for comparisons of fractional equal versus each of the remaining 11 schemas, *within* each population.

## 4.0 DISCUSSION

AuthormetriX (https://sadeosun-a-uthormetrix-v1.streamlit.app/) is designed to facilitate accessibility and adoption of the non-inflationary author credit schemas to address the flaws of the whole count method. Despite the validity and fairness advantage over the whole count schema, the complexities in these schemas have hitherto impeded their adoption and widespread use in administrative processes (e.g., faculty evaluation for tenure and promotion), science policy evaluation, and in bibliometric studies. This suggests that the ease of use of the whole count schema trumps the known superiority of the alternative counting schemas. AuthormetriX therefore levels the playing field by eliminating the complexity barrier which has impeded the adoption of these schemas. With AuthormetriX, users only need to supply the Scopus ID(s) of the author(s)-of-interest, and the relevant corpus of publications downloaded from Scopus. The application then abstracts away the complexities, and time- and labor-intensive calculations of the sequence-dependent author credits, and the summation of those credits for each author-of-interest across all the authors’ publications in the supplied corpus. This is done simultaneously for the whole count and 14 different non-inflationary counting schemas (NIACAS). Given the relationship between collaboration and productivity,[63,64] the output also includes several collaboration metrics.

The use case demonstrates how a researcher’s rank differs based on the counting methods used.[15,18,30,44,45] and the varying degrees of rank displacements with each NIACAS versus the inflationary whole count method (Fig 1a). Given that the fractional equal schema addresses only inflationary bias,[15,18] the schemas detected the bidirectional equalizing bias in ACP to different degrees within each population (Fig. 1b). The schemas were less sensitive to equalizing bias in HCR Clinical Medicine compared to HHI Pharmacy Practice. For both populations, most schemas that emphasize the first author, especially at the expense of the last author (e.g., golden share and geometric standard) detected positive equalizing bias while the ones that emphasize last authors, with or without first-author emphasis (e.g., fractional LAE, fractional FLAE) detected negative equalizing bias. The observation that the authors earned relatively more credit with schemas that emphasize last authors initially suggests that these authors were mostly last authors on their publications. However, the average first or last author proportions (FLAP) were 0.24 and 0.48 for HCR Clinical Medicine and HHI Pharmacy Practice, respectively (Table 2). This means that for HCR Clinical Medicine and HHI Pharmacy Practice respectively, the individuals were neither the first, nor last authors in 76% and 52% of their publications. Thus, they were middle authors who often have only secondary contributions and limited responsibilities on most of their publications.[22–24] Of note, these FLAP values are relatively close to the 0.31 reported for a sample of Highly Cited Researchers (2018 edition) across 5 countries, based on their 2006-2016 publications.[56] Taken together with collaboration metrics in Table 2, the ACP results (Fig. 1b) are consistent with Hagen[15] who concluded that inflationary bias benefits authors with many coauthors and small contributions versus authors with few coauthors. The [standard] harmonic schema detected a negligible equalizing bias based on publication credits in the population studied by Hagen,[18] but that schema detected a significant negative bias in both populations (to different degrees) in the current study. This suggests that the equalizing bias detectable by any given schema would vary by disciplines or populations. This further supports the need for establishing the most appropriate schemas for each discipline.[20,44]

The new harmonic parabolic model (and its formula) reported for the first time in the current study has a few nuances that may make it more accurate for the appropriate discipline. For author counts 5 and above, it rewards the first and last authors more than the arithmetic V, and the percentage extra credit increases with increasing author counts, from 2% to 115%, and 241% for author counts 5, 25, and 50, respectively (see https://sadeosun-a-uthormetrix-v1.streamlit.app/Schema_modeler). Future studies should compare the accuracies of these two schemas relative to empirical author credit data from a relevant discipline.

This example use case demonstrates the extensive bibliometric insights that are possible with AuthormetriX, with little to no *manual* computational efforts and with results obtained within few seconds.

## 4.1 Limitations

There are a few limitations in the use of AuthormetriX. First, the codes are limited to the format of the corpus obtainable from Scopus. Thus, publications in journals not indexed in Scopus would not be included in the analysis. This should be put into consideration especially for disciplines with journals that are not fully indexed in Scopus. However, future updates may include support for corpus obtained from Web of Science. Secondly, Streamlit currently has a 200MB file size limit for single file inputs. This would not be an issue in most cases as the file size of a corpus of 1000 documents is approximately 1.5MB. However, for a much larger corpus, if necessary, the corpus could be limited to columns required in the codes (‘Author(s) ID’, ‘Document Type’, ‘Year’, ‘Title’, and ‘Source title’) to reduce the file size. Thirdly, AuthormetriX leverages the ease and exactness of identifying authors by SCIDs rather than by names that often need disambiguation. However, even with the SCIDs-based search, split, or multiple author profiles are still possible.[65] Thus, the accuracy of the corpus and results depend on identifying an author’s correct profile and unique SCID. Fourthly, the schemas covered in AuthormetriX may not be exhaustive; therefore, when new schemas that meet the inclusion criteria are proposed or discovered, the application would be updated.

Other limitations are those inherited from the nature of the schemas. First, AuthormetriX does not account for shared contributorship (joint first-authorship or joint corresponding-authorship), which has been on the rise.[66] There is also the assumption that the last author who may get a significant fraction of the credit may not necessarily be the senior/corresponding author.[9,27] Finally, AuthormetriX does not address the logical adjustments for first-author corresponding authorship, where both the first and last author credits should belong to only the first author.

## 4.2 Conclusions and Future Directions

AuthormetriX simplifies the implementation of non-inflationary author credit allocation schemas, thus leveraging the fairness, validity, and meritocracy advantages that they have over the whole count method, with minimal cost of time and labor. In addition to opening a new frontier in bibliometric studies, it would be very useful in evaluations of individuals’ scholarly output, for example, in tenure and promotion, awards, grant application and renewal, and rankings. There is no universally valid schema for all disciplines since authorship practices vary among disciplines,[40] and “*one particular accrediting method does not contain the absolute truth*,”[44] therefore, AuthormetriX will facilitate the process of identifying and verifying the appropriate NIACAS for different disciplines. To further lower the barrier of access and use of the NIACAS covered in AuthormetriX, Scopus could integrate the calculations directly into its database. AuthormetriX could usher in the changes in authorship practices that Derek de Solla Price and others had hoped that non-inflationary credit allocation would bring. It could also be the missing link in the yet-to-be-done experiment to test whether the complexity in implementing these schemas was the reason why they have not gained widespread adoption and application. This is relevant because as of this year (*2025)*, valid concerns about the flaws and lack of fairness in the whole count schema were raised over half a century ago.[12]

