## Supplementary material ST1 for "AuthormetriX: Automated Calculation of Individual Authors’ Non-Inflationary Credit-Allocation Schemas’ and Collaboration Metrics from a Scopus Corpus"

Samuel O. Adeosun, RPh, PhD

Supplementary materials

| **Schemas** | **HCR Clinical Medicine** | **HHI Pharmacy Practice** |
| --- | --- | --- |
| fractional equal | 15.8 (12.4-19.2) | 6.0 (4.4-7.7) |
| fractional LAE | 20.0 (15.8-24.2) | 9.1 (6.4-11.7) |
| fractional FAE | 13.3 (10.5-16.0) | 5.0 (3.5-6.5) |
| fractional FLAE | 17.2 (13.6-20.8) | 7.8 (5.5-10) |
| arithmetic standard | 13.2 (10.3-16.2) | 4.5 (3.2-5.9) |
| arithmetic V | 16.7 (13.1-20.3) | 6.8 (5.0-8.7) |
| golden share | 11.4 (8.4-14.4) | 3.8 (2.4-5.3) |
| geometric standard | 11.9 (8.9-15.0) | 4.0 (2.6-5.4) |
| geometric adaptive | 13.2 (10.2-16.2) | 4.6 (3.2-5.9) |
| harmonic standard | 13.1 (10.1-16.0) | 4.6 (3.2-6.0) |
| harmonic parabolic | 16.9 (13.3-20.4) | 6.9 (5.0-8.8) |
| harmonic FLAE | 16.2 (12.8-19.6) | 7.0 (5.1-9.0) |

**Table ST1. Raw Non-Inflationary Author Credit-Allocation Schema Metrics from AuthormetriX**

All pairs (HCR Clinical Medicine vs. HHI Pharmacy Practice) within each row (schema) are statistically significantly different; *p*<0.0001. No *p*-value adjustment for multiple comparisons.
